# Structural insight into CD93 recognition by IGFBP7

**DOI:** 10.1101/2023.06.07.543655

**Authors:** Yueming Xu, Yi Sun, Yuwen Zhu, Gaojie Song

## Abstract

The CD93/IGFBP7 axis are key factors expressed in endothelial cells (EC) that mediate EC angiogenesis and migration. Upregulation of them contributes to tumor vascular abnormality and blockade of this interaction promotes a favorable tumor microenvironment for therapeutic interventions. However, how these two proteins associated to each other remains unclear. In this study, we solved the human CD93–IGFBP7 complex structure to elucidate the interaction between the EGF_1_ domain of CD93 and the IB domain of IGFBP7. Mutagenesis studies confirmed the binding interactions and specificities. Cellular and mouse tumor studies demonstrated the physiological relevance of the CD93–IGFBP7 interaction in EC angiogenesis. Our study provides hints for development of therapeutic agents to precisely disrupt unwanted CD93–IGFBP7 signaling in the tumor microenvironment. Additionally, analysis of the CD93 full-length architecture provides insights into how CD93 protrudes on the cell surface and forms a flexible platform for binding to IGFBP7 and other ligands.

## INTRODUCTION

CD93 (Cluster of Differentiation 93) is a member of group XIV in the C-type lectin-like domain (CTLD) superfamily which is involved in cell adhesion and cytoskeletal organization (*1*). CD93 is highly expressed by endothelial cells (ECs), and there is increasing evidence showing that it plays a key role in their regulation and vessel formation (*2-4*). For example, ECs deficient of CD93 were shown to disrupt actin cytoskeleton organization and reduce adhesion, migration, and tube formation (*5-9*). The adhesion or migration function of CD93 is believed to be facilitated by the association with moesin/cytoskeleton through its intercellular region (*10*). Remarkably, CD93 has been demonstrated to be an attractive target for the treatment of cancer as well as diseases associated with outgrowth of blood vessels (*11-13*). The elevated expression of CD93 in tumor blood vessels are correlated with reduced survival rate in patients with glioblastoma (*14*). Consistently, CD93 signaling contributes to abnormal tumor vasculature, and in multiple mouse tumor models CD93 blockade by monoclonal antibodies promotes tumoral vascular maturation and triggers a substantial increase in intratumoral effector T cells (*15*).

CD93 contains a conserved CTLD, a sushi-like domain (named X), five tandem EGF-like domains, and a mucin-like domain at its extracellular domain (ECD) (*16, 17*). Multiple proteins or molecules have been reported to be the ligands for CD93, which could contribute to the diverse functions for CD93. Not much is known about how CD93 interacts with some of its ligand, such as the glycoproteins dystroglycan, or IL-17D (*5, 8, 18*). Multimerin-2 (MMRN2), an extracellular matrix (ECM) protein that broadly interacts with group XIV CTLD members, engages CD93 through the CTLD domain of CD93 (*9, 12*). Insulin-like growth factor binding protein 7 (IGFBP7), another ECM ligand for CD93, appears to bind to the first two EGF domains of CD93 and does not compete with MMRN2 for CD93 binding(*15*). IGFBP7 protein contains an IGF-binding (IB) domain at its N terminus, a Kazal-type serine proteinase inhibitor domain (Kazal) in its central region, and an Ig-like C2-type (IgC2) domain at its C terminus (*19*). IGFBP7 interacts with CD93 via its IB domain, however, the exact interactions between these two molecules are yet to be determined (*15*).

Here, we report on the CD93–IGFBP7 complex structure, which reveals molecular details between key domains of these two proteins. Mutagenesis studies on molecular and animal levels confirm the binding specificities and physiological relevance of the CD93–IGFBP7 interaction. Furthermore, analyzation of CD93 full-length architecture lays a basis for how CD93 protrudes on the cell surface and forms a flexible platform for its binding to IGFBP7 and other ligands.

## RESULTS

### Structural determination

Our previous work has identified the EGF_1,2_ domains on CD93 and the IB domain on IGFBP7 as key determinants for the complex formation (*15*). To form a stable complex between the two proteins, we first individually expressed different fragments containing the key domains for each protein. However, 1:1 mixture of fragments from IGFBP7 and CD93 yielded neither a stable complex upon gel filtration chromatography nor crystal hit upon screening, potentially as a result of a relatively low affinity between the two proteins. Learning from our previous experience in the successfully structural determination of the CD97–CD55 complex (*20*), we employed a 12-residue linker to connect the C-terminus of IGFBP7 fragment composed of the IB-Kazal domains (henceforth termed IBK) with the N-terminus of CD93 fragment composed of the CTLD-X-EGF_1,2_ domains (henceforth termed CXE_12_). This chimeric protein was successfully expressed and purified from HEK293 cells (Method). Homogeneity of the chimeric protein was verified by size exclusion chromatography with multiangle light scattering (SEC-MALS), which indicated a monomeric state for the chimeric protein (fig. S1A). We then successfully crystallized the chimeric protein to 3.24 Å resolution. The structure was solved with molecular replacement using the AlphaFold2 models as templates (*21, 22*), and the final structure was refined to R_work_ and R_free_ of 0.244 and 0.302, respectively (Table S1).

### Overall structure

There are two molecules for each CD93 and IGFBP7 fragment in the final structural solution, with molecules B and C showing slightly improved density compared to the other two molecules (Fig. 1A). The linker region is not modelled because of its intrinsically disorder nature. Similarly, densities are also missing for the residues 227-231 of CD93 and residues 91-103 of IGFBP7. Of note, in the final model, the whole EGF_2_ domain was not resolved in either CD93 molecule, despite its presence in the crystals as shown by SDS-PAGE (fig. S1B). The crystal packing suggests that the EGF_2_ must be disordered since in the lattice the C-terminus of EGF_1_ domain lies adjacent to a large void that may accommodate multiple orientations of the EGF_2_ domain (fig. S1C).

**Fig. 1.**
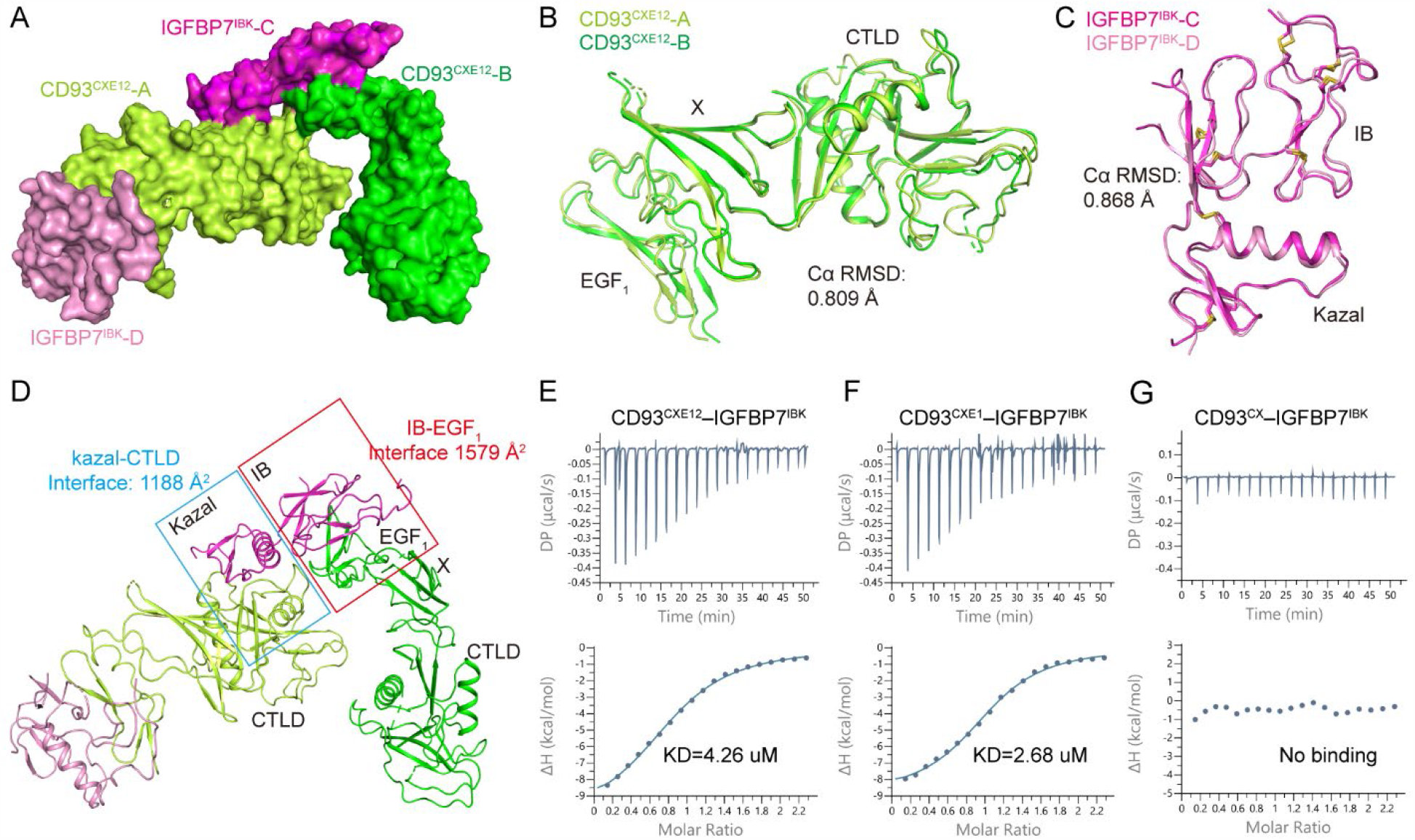
CD93–IGFBP7 overall structure. A. The asymmetric unit in the CD93–IGFBP7 complex structure. B-C. Superposition of the CD93 chains (B) and IGFBP7 chains (C). D. The two main CD93–IGFBP7 interfaces revealed in the crystal lattice. The Kazal–CTLD and IB–EGF_1_ interfaces are framed and labelled. In A-D, the CD93 and IGFBP7 chains are color coded with their labels. E-G. Binding affinities of IGFBP7 with the CD93 fragments. CXE12, CXE1, and CX stand for CTLD-X-EGF_12_, CTLD-X-EGF_1_, CTLD-X fragments, respectively; IBK, IB-Kazal fragment.

The two CD93 molecules in the asymmetric unit align to each other quite well, as do the two IGFBP7 molecules (Fig.1B-C). Remarkably, the comparison of CD93 molecules show essentially identical orientations between the tandem CTLD, X, and EGF_1_ domains (Fig. 1B). This structural feature is secured by extensive contacts between the adjacent domains (see below for details). For the EGF_1_ domain of CD93, on one side it forms stable contacts with the X domain through its loop regions, whereas on the opposite side it packs against the IB domain of IGFBP7 in the lattice. The IB domain of IGFBP7 features a ladder-like shape and the all-sheet structure is facilitated by 6 pairs of disulfide bonds and extensive intradomain hydrogen bonds (Fig. 1C). The IGFBP7 IB domain forms extensive contacts with the Kazal domain through a key interdomain disulfide bond (C113– C131) as well as several hydrophilic interactions.

### The CD93–IGFBP7 binding mode

The lattice packing of CD93–IGFBP7 suggest two potential interfaces between the proteins: one between the IB domain of IGFBP7 and the EGF_1_ of CD93; the other between the Kazal domain of IGFBP7 and the CTLD domain of CD93 (Fig. 1D). To validate which interface is physiological relevant, we expressed different fragments of CD93 and measured their binding affinities with the IBK fragment of IGFBP7 by isothermal titration calorimetry (ITC). The ITC titrations showed that the fragments of IBK and CXE_12_ can bind to each other with a KD of 4.26 μM (Fig. 1E). Of note, the shorter fragment of CXE_1_ (with EGF_2_ removed) can bind to IGFBP7 with similar level of affinity (2.68 μM) (Fig. 1F), in contrast to a total loss of binding when the EGF_1_ domain was further removed (Fig. 1G). These results, together with the single point mutation data shown below, strongly suggest that the binding between CD93 and IGFBP7 relies on the IB–EGF_1_ interface. Thus, we subsequently focused on this interface for further analysis.

The ladder-like IB domain of IGFBP7 binds through one side to the CD93 EGF_1_ domain and the interaction buries in total 1579 Å^2^ of solvent-accessible area (Fig. 2A). The IB–EGF_1_ interface can be divided into two parts: on one side, the E^52^TR^54^ tripeptide of the IB domain forms extensive hydrophilic interactions with the CD93 EGF_1_ domain residues R303, C285, L293, and D295; on the other side, the Y^80^CAP^83^ tetrapeptide of IB domain forms a hydrophobic patch jointly with L50 and A64 that covers the bulky F276 of EGF_1_ on CD93 (Fig. 2B, fig. S2). The conformation of the tetrapeptide is stabilized by an intradomain disulfide bond (C81–C111) within the IB domain. In addition, Y80 also hydrogen bonds to the S282 of EGF_1_. Fig. 2C shows how the corresponding domains of the two proteins can accommodate each other in terms of both physical shape and electrostatic potential. For example, the E52 and R54 of IGFBP7 point to a basic and an acidic region on the surface of CD93, respectively. Also, the hydrophobic residues of IGFBP7 (A64, Y80, and P84) are placed in a concave and hydrophobic surface on CD93, thus avoiding steric clash and forming hydrophobic interactions that stabilize the complex.

**Fig. 2.**
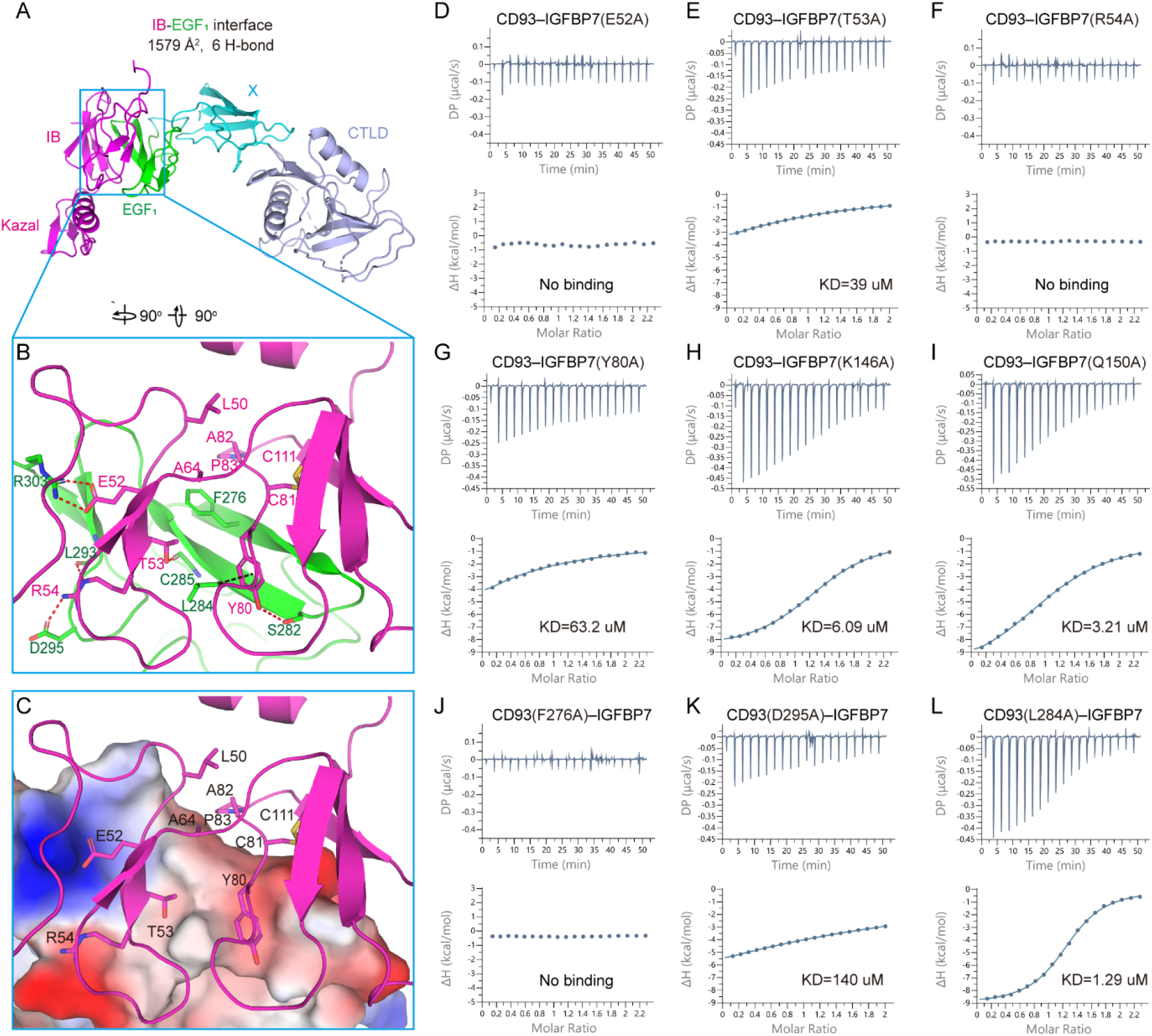
Detailed interactions of CD93–IGFBP7. A. Overall structure of the CD93–IGFBP7 complex. Domains of CD93 and IGFBP7 are color coded. B. Close-up view of the interface between the CD93 EGF_1_ and IGFBP7 IB domains. Key residues involved in interactions are shown as sticks and are labeled. Red dashed lines indicate hydrogen bonds. Black dashed line indicates van de Waals interaction between L284 (Cβ) of CD93 and Y80 of IGFBP7. C. Same view as (B) with the CD93 shown as electrostatic potential. D-I. Binding affinities of WT CD93^CXE12^ with IGFBP7^IBK^ mutants. J-L. Binding affinities of CD93^CXE12^ mutants (F276A, D295A, and L284A) with WT IGFBP7^IBK^. CXE12 and IBK stand for CTLD-X-EGF_12_domains of CD93 and IB-Kazal domains of IGFBP7, respectively.

To validate the characterized interactions between the two domains, we conducted single point mutations on both sides of the complex and measured the binding affinities (Fig. 2D-L). The T53A and Y80A mutations on IB domain decreased their binding affinities by ∼9- and >10-fold, respectively (Fig. 2E, G). Remarkably, the E52A and R54A mutants of the IB domain each produced an unmeasurable binding with CD93 (Fig. 2D, F), consistent with the fact that they form at least one salt bridge with the counterparts in CD93. As negative controls, the Kazal domain mutations on the Kazal–CTLD packing interface (K146A, Q150A) did not significantly affect the binding with CD93 (Fig. 2H, I). On the other side, the F276A mutant of EGF_1_ also totally abolished binding with the IB domain of IGFBP7, suggesting the hydrophobic interactions contributed by F276 are of vital importance for complex formation (Fig. 2J). The R54 interacting residue, D295, if mutated, reduces the binding affinity by ∼33-fold (Fig. 2K). These ITC results align well with our structural findings and further support the physiological importance of the characterized IB– EGF_1_ interface.

The E^52^TR^54^ tripeptide of IGFBP7 and its counterparts in CD93 are highly conserved among different species. In contrast, the tyrosine in the tetrapeptide of IGFBP7 is only seen in *human* and *callithrix*, and in other species the corresponding position of Y80 is a histidine (fig. S3). Interestingly, in the opposing region in CD93, the hydrophobic L284 is also only conserved in *human* and *callithrix*, while in other species it is replaced by an arginine (fig. S4). L284 of CD93 forms a van de Waals interaction with Y80 of IGFBP7 but mainly through its Cβ atom (Fig. 2B). This is probably the reason why the L284A mutation did not significant compromise its affinity with IGFBP7 (Fig. 2L). Nevertheless, the histidine and arginine are more hydrophilic, and they may hydrogen bonding to each other. Hence, the paired presence of Y/L and H/R in these species suggests how the receptors are evolved to accommodate the environmental change in their ligands, or vice versa.

### The CD93–IGFBP7 interaction in EC angiogenesis

Based on the CD93–IGFBP7 complex structure, we further performed mutagenesis to study the impact of this interaction on EC functions. CD93 and IGFBP7 are constitutively expressed in HUVEC cells and mediate EC angiogenesis and migration(*9, 15, 23*). We knocked out CD93 in HUVEC cells by the CRISPR/Cas9 technology and the loss of surface CD93 expression was confirmed by flow cytometry. We sorted out CD93KO cells and performed an EC-mediated wound healing assay. CD93KO HUVEC cells were significantly slower in migration than its WT counterpart (Fig. 3A), which is consistent with previous publications(*5, 14*). We transduced CD93KO HUVEC cells with different CD93 molecules to assess their capacity of promoting EC migration. Transduction of WT CD93 was able to improve EC migration in CD93KO HUVEC cells, which became comparable to WT HUVEC cells; however, CD93 mutants that lose the IGFBP7 binding, including F276A, D295A, and double mutant F276A/D295A, failed to do so (Fig. 3A).

**Fig. 3.**
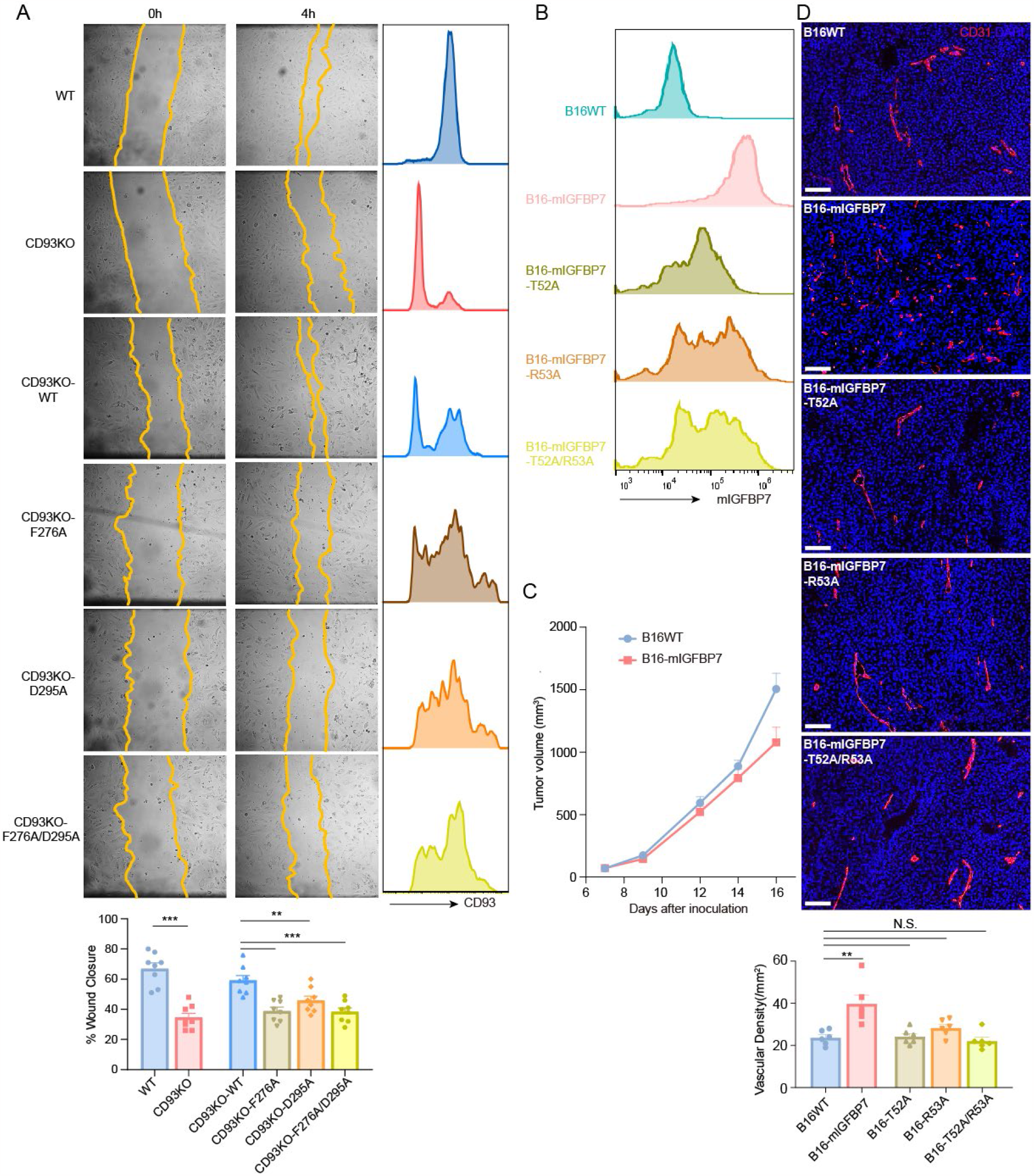
Mutagenesis study of the CD93–IGFBP7 interaction in EC functions. A. HUVEC cells were knocked out of CD93 via the CRISPR/Cas9 technology and loss of surface CD93 was confirmed by flow cytometry (right). CD93KO HUVEC cells were transduced to express WT CD93 or different CD93 mutants. Their migratory ability was assessed in a wound-healing assay. B-D. B16F10 tumor cells were transduced to express different mouse IGFBP7 mutants. Their expression in B16F10 tumor cells were examined by intracellular staining (B). B16 WT and B16 tumors transduced to express WT IGFBP7 at the number were subcutaneously inoculated into WT B6 mice. Tumor growth curves was recorded (C). The densities of CD31^+^ tumor blood vessels were assessed by immunofluorescent staining (D).

We used a mouse tumor model to study the impact of IGFBP7 binding to CD93 on tumor angiogenesis. B16F10 melanoma cells do not secrete or produce IGFBP7, based on intracellular staining (Fig. 3B). We transduced B16F10 with WT IGFBP7 and subcutaneously implanted them into black B6 mice. Although IGFBP7 expression in B16 tumors did not affect tumor growth *in vivo* (Fig. 3C), B16F10-IGFBP7 tumors contained significantly more CD31^+^ blood vessels, compared to control B16F10 tumors (Fig. 3D). This implies that IGFBP7 produced by tumor cells is able to promote vascular angiogenesis. We also transduced B16 tumors to express different mouse IGFBP7 mutants that presumably lose the binding to CD93, including T52A (corresponds to human T53A), R53A (corresponds to human R54A), and double mutant T52A/R53A. Analysis of tumor tissues revealed a similar extent of vascular densities between these mutant transducers and control B16 tumors (Fig. 3D). Taken together, our studies supported an indispensable role for the IGFBP7–CD93 interaction in EC angiogenesis.

### Specializations for IGFBP7

IGFBP7 is quite unique among the totally seven members within the IGFBP7 family. The other six members each contain an N-terminal IB domain that connects via a disordered linker to a C-terminal IB domain, and the two IB domains work cooperatively for their high-affinity IGF binding (Fig. 4A)(*24*). For IGFBP7, the C-terminal IB domain is missing; instead, it includes an Ig-like domain at the C-terminus and a Kazal domain in the middle (Fig. 4B). Structural and sequence comparisons suggest that the N-terminal IB domains of IGFBP1-6 bind to IGF1 through the tip region, whereas the homologous domain in IGFBP7 binds through one side to its ligand CD93 (Fig. 4C-D). The key residues in IGFBP1-6 for IGF1-binding are not conserved in IGFBP7 (fig. S5). Furthermore, IGFBP1-6 each contain two hydrophobic residues at the N-terminus that bound and covered a non-polar region in the IGF1, an important region for binding with IGF1R (Fig. 4C). In contrast, the IGFBP7 is rather hydrophilic in its N-terminus (fig. S5). These structural features explain why IGFBP7 does not bind to IGFs with similar affinity compared to other members(*24, 25*).

**Fig. 4.**
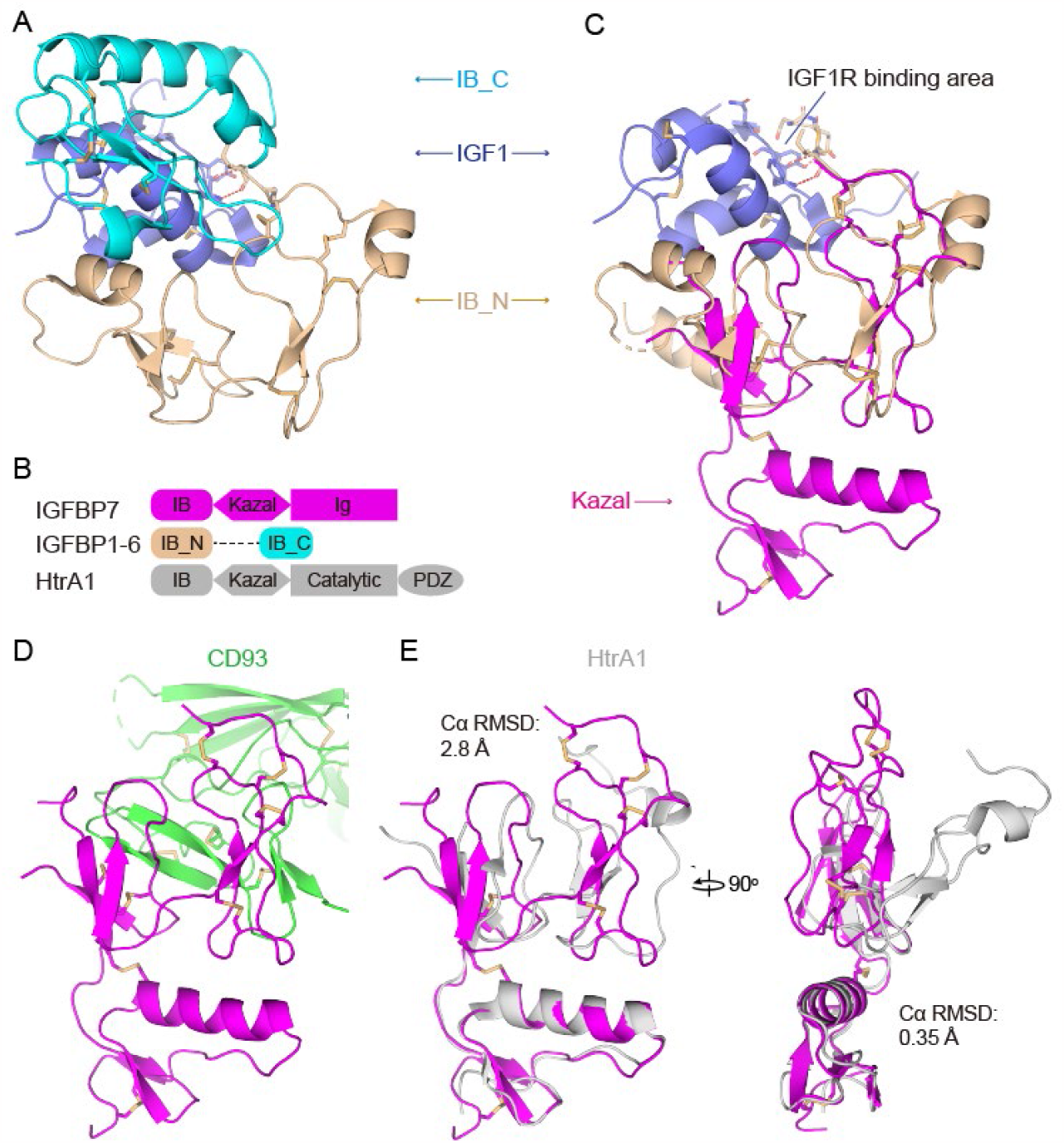
Specializations of IGFBP7. A. The complex structure of IGFBP4–IGF1 (PDB ID: 2DSR). Both N-terminal and C-terminal IB domains of IGFBP4 are involved in the interaction with IGF1. B. Schematic diagrams represent the structure of IGFBP7, other IGFBP members (IGFBP1-6), and HtrA1. C. Superposition of IGFBP7 onto the IGFBP4 (IB_N)–IGF1complex (PDB ID: 2DSP). D. IGFBP7–CD93 binding mode. E. Superposition of IGFBP7 and HtrA1 on the IB (left) and Kazal (right) domains. The HtrA1 structure is from PDB ID 3TJQ. All the structures are color coded with their labels.

A Dali (*26*) search against IGFBP7 revealed the closest structure of IGFBP7 is the N-terminal IB-Kazal modules of HtrA1(*27*), which shares a Cα r.m.s.d of 2.8 Å for the IB domain and 0.35 Å for the Kazal domain (Fig. 4B, E). Moreover, the overall shape of the IGFBP7 IB domain looks more like a flat rectangle, in contrast to a somewhat distorted shape in the counterpart of HtrA1 (Fig. 4E), which does not support a similar binding mode as in the CD93–IGFBP7 complex. Most importantly, the key CD93-binding region, E^52^TR^54^ tripeptide, is not conserved in other IGFBP members, as well as in other IB domain-containing proteins (*e*.*g*., IGFBPL1, Kazal D1 and HtrA1).

This analysis suggests that, among these homologous proteins, IGFBP7 is the only ligand that can bind to CD93, thus exclusively triggers CD93-directed physiological consequences.

### Domain organization of CD93

The structure reveals that the tandem CTLD, X, and EGF_1_ domains formed extensive contacts between adjacent domains, which buried > 1000 Å^2^ of surface area for each interface (Fig. 5A). The X–EGF_1_ interface is mostly hydrophobic, composed of two loops in each domain that are rich in glycine and proline residues (Fig. 5B). At the edge of the interface, P196 of the X domain forms a hydrophobic triad core with the EGF_1_ domain residues Y262 and F266. At the middle, P255 and V258 of the X domain are packed against the C271–C285 disulfide and the F283 and V298 of EGF_1_ domain. In addition, there are some polar interactions within the X–EGF_1_ interface, including a hydrogen bond contributed by N267 and the main-chain of L193. The CTLD–X interface is also mainly hydrophobic in the core, with residues V29, C31, C36, T38, F182 and F184 of the CTLD domain and residues M187, P204, F205, L212 and P216 of the X domain participating (Fig. 5C). This hydrophobic core is flanked by several pairs of hydrogen bonds contributed by CTLD residues E27, C31, N56, and R74, and the X domain residues T207, A214, Q206, and E213. Most of these non-polar residues are highly conserved within the CD93 orthologs (Extended Data Fig. 3). It appears that the X and EGF_1_ domains each turn and fold up against their precedent domain to cover these hydrophobic residues on each domain.

**Fig. 5.**
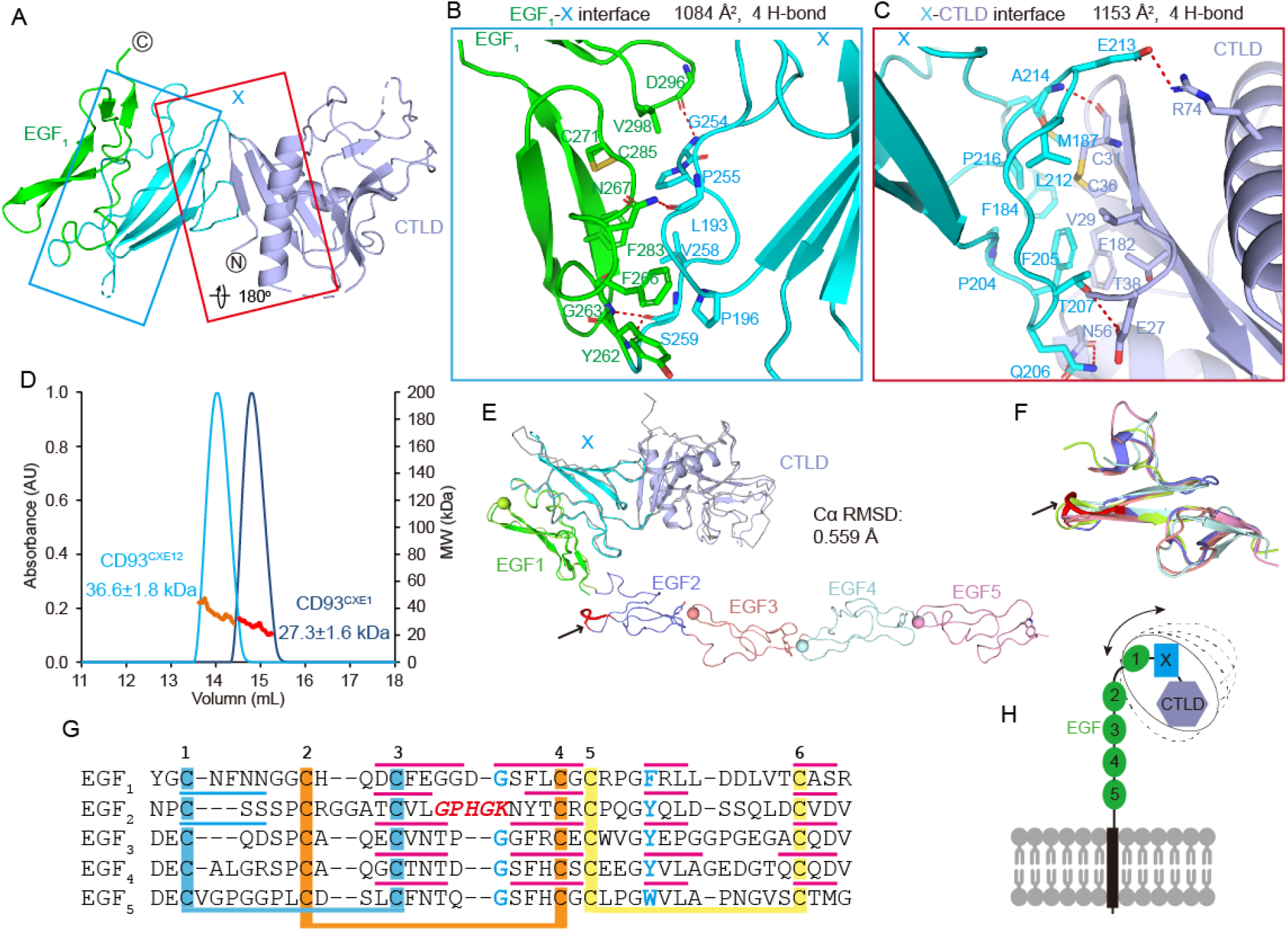
Domain organization of CD93. A. Overall structure of CD93. CD93 domains are colored as in Fig. 2A. X–CTLD and EGF_1_–X interfaces are marked with red and teal boxes, respectively. B-C. Detailed interactions of the CD93 EGF_1_–X (B) and X–CTLD (C) interfaces. Key residues involved in interactions are shown as sticks and are labeled. Red dashed lines indicate hydrogen bonds. D. SEC-MALS profiles of CD93^CXE12^ and CD93^CXE1^. E. Superposition of the CD93^CXE12^ crystal structure with the predicted full-length CD93 model from AlphaFold2. Domains within the CD93^CXE12^ crystal structure are colored as in A. For clarity, the corresponding domains in the predicted model are colored gray, and the remaining EGF_2-5_ domains are color coded. Key interaction residues between EGF domains are shown as spheres (glycines) or sticks (aromatic residues) and are highlighted in G. F. Superposition of EGF domains from the AlphaFold2 CD93 model. G. Sequence alignment of EGF domains (EGF_1-5_) in CD93 AlphaFold2 model. Horizontal cyan and red thin lines indicate α-helices and β-strands, respectively. Disulfide bonded cysteines are linked with thick lines. In E-G, the loop bearing GPHGK is in bold and colored red. H. Full-length CD93 model on the cell surface. Dashed ellipses and arrow indicate the flexibility of the N-terminal domains.

The CTLD–X interface revealed in our complex structure is almost identical to the recently solved CTLD-X fragment crystal structure (*28*). The CTLD-X fragment structure also indicated an anti-parallel dimeric conformation by the lattice packing(*28*). In fact, our crystal structure shows similar packing between CD93^CXE12^-A and its symmetry-related molecule, whereas the packing is totally different for CD93^CXE12^-B and there is no potential dimeric interface around CD93^CXE12^-B. Hence, to further validate the conformational state of CD93 in solution, we measured the molecule mass of different fragments by SEC-MALS. For CD93 fragments CXE_1_ and CXE_12_, the measured molecular mass of 27.3±1.6 kDa and 36.6±1.8 kDa are close to the calculated molecular mass of 31 kDa and 35.4 kDa, respectively (Fig. 5D). Thus, these results suggest that the CD93 extracellular region is a monomer, and the dimeric conformation visualized in the lattices is likely an artifact. Although we cannot exclude the possibility that full-length CD93 may form a concentration-dependent dimer on the cell surface, the binding interaction between CD93 and IGFBP7 may not be affected by the potential dimer mediated by the CTLD-X region.

The CTLD-X-EGF_1_ domain organization was surprisingly predicted by AlphaFold2 although with subtle vibrations (Fig. 5E), suggesting the AlphaFold2 models can be a good template or complement for future structural and functional investigation. AlphaFold2 also predicted a distinct orientation between the EGF_1_ and EGF_2_ domains compared to a conserved orientation between the subsequent EGF_2-5_ domains (Fig. 5E). The tandem EGF domains are frequently seen in adhesion proteins, and, in many cases, the adjacent EGF domains form steady orientations that are facilitated by typical interactions between an aromatic residue in the precedent domains and a glycine residue in the following domains(*20*). The aromatic and glycine residues are highly conserved within the CD93 EGF domains except EGF_2_. Specifically, in EGF_2_ the loop bearing the conserved glycine residues is longer, and the inserted bulky H322 and K324 likely distorts the loop and blocked the interaction with the preceding EGF_1_ domain (Fig. 5F-G). Hence, we reasoned that the disordered EGF_2_ domain is a consequence of its flexible orientation since the consensus interaction is not applicable between the EGF_1_ and EGF_2_ domains of CD93. Also, the flexibility between the EGF_1_ and EGF_2-5_ domains may provide a chance for CD93 to smoothly present its N-terminal CTLD through EGF_1_ domains for the binding with its ligands (Fig. 5H).

## DISCUSSION

Generally, IGFBP family members 1-6 function by binding IGFs (insulin/IGF1/IGF2) thus regulating their availability in body fluids and tissues and modulating IGFs’ binding to their receptors, which are essential for body growth, metabolism and survival(*29, 30*). Nevertheless, IGFBP7 binds IGFs with much lower affinity, and hence was suggested to function mainly through IGF-independent manners (*19*). Here we determined the complex structure of CD93–IGFBP7 in atomic resolution, by which we revealed the specializations for IGFBP7 compared to IGFBPs 1-6 and structurally explained the low affinity binding of IGFBP7 with IGFs. IGFBP7 is also involved in cell adhesion by binding to αvβ3 (*23*), type IV collagen (*31*), and heparin sulfate (*32*). The determined structure of IGFBP7 provided a basis for further study of IGFBP7’s interactions with these ligands and their functional consequences.

Our analysis on the CD93 fragment structure and its full-length architecture also provided a template for future investigation for this function. Beside IGFBP7, another ligand Multimerin-2 has been suggested to bind to the X domain of CD93, through mainly the F238 within the X domain (*8*). The structure of CD93 reveals that F238 is located in the middle of the X domain and its side-chain is half exposed. Interruption of the interaction between CD93 and MMRN2 has been suggested to induce disruption of vascular integrity in tumors, representing a new target for pharmaceutical intervention (*14*). The hydrophobic environment around F238 indicated that the binding between CD93 and Multimerin-2 is mainly through van de Waals interactions. Although F238 is also located adjacent to the IGFBP7-binding region, the EGF_1_ domain, the binding of CD93 with the two ligands was proven not to be interfered by each other (*15*). Our mutagenesis study also found an indispensable role of the CD93–IGFBP7 interaction in EC angiogenesis, suggesting that MMRN2 and IGFBP7 play a non-redundant role in CD93-mediated EC function.

The blockade of CD93–IGFBP7 interaction has been shown to be potentially beneficial for the treatment of cancer, especially when in combination with anti-PD therapy(*15*). In addition, IGFBP7 was found markedly upregulated in tumor blood vessels and capable of promoting vascular angiogenesis (*33*). The CD93–IGFBP7 complex structure reported here shed lights on the binding determinants between these two proteins. Mutagenesis studies confirms the binding interface and explains the binding specificities between CD93 and IGFBP7, as well as the involvement of this interaction in vascular angiogenesis. The current study lays a basis for potential development of small molecular blockade to precisely break the excessive CD93–IGFBP7 interaction in the tumor microenvironment.

## METERIALS AND METHODS

### Molecular cloning

Codon-optimized cDNA sequence of human CD93 and IGFBP7 were synthetized by TONGYONG (Anhui, China). A CD93 CTLD-X-EGF_12_ fragment, a IGFBP7 IB-Kazal fragment, and a chimeric complex were constructed on a customed pLEXm(*34*) vector with N-terminal signal peptide and C-terminal His6 tags. In the chimeric construct, a 12-residue flexible linker was added to link the C terminus of the IGFBP7 fragment and the N terminus of the CD93 fragment (named IGFBP7–12a–CD93). Truncations and mutations of CD93 were generated by splicing overlap extension (SOE) PCR using TransStart Fast-Pfu DNA polymerase (TransGen, China) and cloned to the same vector for expression and purification. All constructs were confirmed using DNA sequencing.

### Protein production and purification

HEK293F GnTI^-^ cells were cultured with FreeStyle 293 Expression Medium (BasalMedia, China) without serum in suspension and maintained in a 90 rounds per minute rotating (50 mm) amplitude shaker, at 37 °C, 5% CO_2_, and 95% humidity. When cultured to 1-2 million/mL, the cells were transiently transfected with DNA: polyethylenimine at 1:3 wt/wt. After 3 to 5 days, cell supernatants were harvested and incubated with Ni-NTA beads (Smart-Lifesciences, China) at 4°C for about 2 h. Beads were washed with 20 mM HEPES, pH 7.5, 500 mM NaCl, and 30 mM imidazole, and eluted with 20 mM HEPES, pH 7.5, 500 mM NaCl, and 300 mM imidazole. The proteins were further purified using size exclusion chromatography (SEC) with a Superdex S200 increase gel filtration column (cytiva) equilibrated with 20 mM HEPES, pH 7.5, and 150 mM NaCl. The chimeric complex IGFBP7–12a–CD93 was concentrated to 7 mg/mL for crystallization trials, while the truncations and mutants of CD93 or IGFBP7 were concentrated to optimal concentration and subjected to ITC or SEC-MALS analysis.

### Crystallization and structure determination

The concentrated IGFBP7–12a–CD93 complex was set up for crystallization using hanging drop with NT8 (Formulatrix). Crystals grew under several conditions, but the highest diffraction crystals were produced at 18 °C in 0.1 M imidazole, 20% w/v PEG monomethyl ether 2000, and 0.2 M ammonium citrate tribasic. Single crystals were directly frozen in liquid nitrogen. Diffraction data were collected at beamline BL10U2 at the Shanghai Synchrotron Radiation Facility. Data were indexed, integrated, and scaled using the automatic XIA2 software package(*35*). The structure was solved by the molecular replacement method using the CD93 AlphaFold2(*21, 22*) model as the search template. Refinement was carried out using Phenix(*36*) and with manual adjustments with Coot(*37*).

### Isothermal titration calorimetry (ITC) assay

ITC experiments were performed using a PEAQ-ITC instrument (Malven) at 25 °C. All protein samples were prepared in assay buffer containing 20 mM HEPES, pH 7.5, and 150 mM NaCl. The assays were performed in a multi-injection mode by titrating 40 μL of titrant in the syringe into 200 μL of titrate in the cell. To characterize the interaction between CD93 and IGFBP7, titrations were conducted with 20 injections of 250 μM CD93 (truncations or mutants) into 25 μM IGFBP7 (WT or mutant) at intervals of 150 s.

### Size exclusion chromatography with multi-angle static light scattering (SEC-MALS)

The SEC-MALS measurements were carried out using the Agilent 1260 infinity II HPLC system coupled to a DAWN HELEOS-II light scattering detector (Wyatt Technology) and an Optilab TrEX refractive index monitor (Wyatt Technology). 100 μL of CD93 fragments CTLD-X-EGF_12_ or CTLD-X-EGF_1_ was loaded onto a Superdex S200 increase gel filtration column (cytiva) equilibrated with buffer 20 mM HEPES, pH 7.5, and 150 mM NaCl at a flow rate of 0.5 mL/min. The SEC-MALS/RI data was collected and analyzed with the ASTRA software (Wyatt Technology).

### HUVEC experiments

Pooled HUVEC cell line was purchased from ThermoFisher (Cat# C01510C). HUVEC cells were cultured in Endothelial Cell Growth Medium from Sigma-Aldrich (#211-500) at 37°C, 5% CO_2_, and 95% humidity. Lipofectamine CRISPRMAX Kit was utilized to generate CD93-knockout (KO) HUVEC cells following the instruction from the manufacturer. Cas9 nuclease and human CD93 multi-guide mod sgRNA (SO# 11538146) were purchased from SYNTHEGO. CD93KO HUVEC cells were enriched by negative selection using MACS magnetic separating system including anti-human CD93 mAb (SinoBiological #12589-MM01), anti-mouse IgG microbeads (Miltenyi Biotec #130-048-401) and MS columns (Miltenyi Biotec #130-042-201). Human CD93 WT, F276A, D295A and F276A/D295A mutants were reconstituted in CD93KO HUVEC cells by lentivirus transduction. The Lentivirus producing system was kindly provided by M Eric Kohler M.D. at CU Anschutz, including Lenti-X cell line, pMDLg/pRRE, pMD2.G and pRSV-Rev co-plasmids. CD93 WT and mutants lentiviral construct DNA were synthesized and purchased from GenScript. After transduction, CD93 expression (including all mutants) was validated by flow cytometry (anti-CD93-APC, BioLegend, #336120). For the wound healing assay, HUVEC cells were seeded into a 24-well plate at 5×10^4^ /well one day in advance. After scratches, images were taken at 0 and 4 hours. The area of the wound was calculated by ImageJ software (Ver. 1.53t).

### Mouse tumor vasculature

B16F10 cells were cultured in RPMI1640 medium with 10% FBS. After the cells were transfected with mouse construct of IGFBP7 WT and mutants (GenScript) and enriched by antibiotic selection, mouse IGFBP7 expression was validated by intracellular IGFBP7 staining (anti-mouse-IGFBP7 clone 2C6). 2×10^5^ B16 tumor cells in 100ul HBSS buffer were prepared and inoculated to right flank of 6-week-old female C57BL/6J (Jax) mice. 12 days after inoculation, tumor tissues were collected for FFPE block and sections. Tissue slides were rehydrated and gone through antigen retrieval for downstream immunofluorescent counterstaining of mouse CD31 (abcam, #ab28364) and nucleus (DAPI, Invitrogen, #D1306). All bright fields and IF images were taken and analyzed by Zeiss Axio Observer microscope with Slidebook 3i software (Intelligent Imaging Innovations, Inc.). All flow cytometry data and plots were analyzed and generated with FlowJo software (Ver. 10.8.1). The bar plots were created by GraphPad Prism (Ver. 8.4.3), as well as the determination of statistical significance (p<0.05).

## Acknowledgments

We thank the staff of BL10U2 from shanghai Synchrotron Radiation Source for their assistance in data collection and processing. We thank the Instruments Sharing Platform of School of Life Sciences, East China Normal University. This work was supported by the National Natural Science Foundation of China (32171215) (to G.S.); the basic research program of Science and Technology Commission of Shanghai Municipality (21JC1402400) (to G.S.); NIH R01CA269644 (to Y.Z.); R01CA258302 (to Y.Z.); R01CA279398 (to Y.Z.).

## Author Contributions

Y.X. optimized constructs, expressed and purified the complex proteins for crystallization, conducted mutagenesis, and edited the initial manuscript; Y.S. conducted HUVEC experiments and Mouse tumor vasculature experiments. Y.Z. supervised the cellular and mouse studies, and edited the manuscript. G. S. supervised the project, analysed the data, and wrote the manuscript. All authors involved in the discussion and provided feedback for the manuscript.

## Data and materials availability

Atomic coordinate and structure factor for the CD93–IGFBP7 complex has been deposited in the Protein Data Bank with identification code 8IVD. Correspondence and requests for materials should be addressed to G.S. or Y.Z.

